# The contribution of PIP2-type aquaporins to photosynthesis in response to increased vapour pressure deficit

**DOI:** 10.1101/847053

**Authors:** D. Israel, S. Khan, C.R. Warren, J.J. Zwiazek, T.M. Robson

## Abstract

Roles of three different plasma membrane aquaporins (PIPs) in leaf-level gas exchange of *Arabidopsis thaliana* were examined using single, double and triple knockout mutants and compared to the Columbia-0 wild type (WT) plants. Since multiple *Arabidopsis* PIPs are implicated in conducting carbon dioxide across membranes, we focused on identifying whether the examined isoforms affect photosynthesis, either mediated through the control of stomatal conductance to water vapour (*g*_s_) or mesophyll conductance of CO_2_ (*g*_m_) or a combination of both. In two separate studies, we grew *Arabidopsis* plants in a low humidity environment and under high humidity conditions. We found that the contribution of functional PIPs to *g*_s_ was larger under conditions of low air humidity when the evaporative demand was high, whereas any effect of lacking PIP function was minimal under higher humidity conditions. The *pip2;4* knockout mutants had 44% higher *g*_s_ than the WT under low humidity conditions, which in turn resulted in an increased photosynthetic rate (*A*_net_). *At*PIP2;4 is thus likely to be involved in maintaining a positive water balance and high water use efficiency through mediation of transmembrane water flow. The lack of functional *AtPIP2;5* on the other hand did not affect *g*_s_, but reduced *g*_m_ indicating a possible role in regulating CO_2_ membrane permeability. This potential regulatory function was indeed confirmed by subsequent stopped flow measurements of yeast expressing *At*PIP2;5.

## Introduction

Water flow across membranes, and thus through the plant, is regulated by aquaporins, which in addition to water may also conduct small neutral molecules and gasses including CO_2_ and O_2_ (Agre et al., 1993; Heckwolf et al., 2011; Zwiazek et al., 2017). *Arabidopsis thaliana* possesses 35 different aquaporin isoforms that are divided into four subfamilies (Johanson et al., 2001): plasma membrane intrinsic proteins (PIPs) located in the plasma membrane (Daniels et al., 1994; Kammerloher et al., 1994; Kaldenhoff et al., 1995; Hachez et al., 2014), tonoplast intrinsic proteins (TIPs) localised to the tonoplast (Maurel et al., 1993; Beebo et al., 2009), Nodulin26-like intrinsic proteins (NIPs) with various membrane locations (Mizutani et al., 2006; Choi and Roberts, 2007), and small basic intrinsic proteins (SIPs) that are found in the membranes of the endoplasmic reticulum (Ishikawa et al., 2005). PIPs have been shown to be involved in a variety of processes regulating plant water flow starting from the root through the stem as well as into and out of the leaves (Javot et al., 2003; Fraysse et al., 2005; Da Ines, 2010; Ben Baaziz et al., 2012; Gambetta et al., 2013). Based on their phylogeny, PIPs are further divided into two subgroups, the PIP1s and PIP2s, with five and eight isoforms respectively (Johanson et al., 2001). Water-permeability varies between the isoforms (Kammerloher et al., 1994; Kaldenhoff et al., 1995; Kaldenhoff et al., 1998; Chaumont et al., 2000; Li et al., 2015) and in fact, PIP1s are believed to transport water only when part of the heterotetramer structure with PIP2s (Fetter et al., 2004; Zelazny et al., 2007; Otto et al., 2010).

Standard greenhouse conditions for plant research provide ample light, water, nutrients and temperature, and thus rates of photosynthesis (*A*_net_) are largely determined by the availability of CO_2_, which is limited by two resistances in series. First by the diffusion of CO_2_ from the leaf exterior into the intercellular airspaces through the stomata and second by the diffusion from intercellular airspaces into the chloroplast, as described by mesophyll conductance (*g*_m_). In *A. thaliana*, *At*PIP1;2 was the first aquaporin identified to be a significant contributor to *g*_m_ due to its CO_2_ permeability (Heckwolf et al., 2011), but *At*PIP1;4 has now also been recognised to facilitate CO_2_ diffusion across plasma membranes (Li et al., 2015). Isoforms of the PIP2 subgroup were believed to be specific to water until they were discovered to also conduct H_2_O_2_ (Dynowski et al., 2008), which is structurally very closely related to H_2_O. However, recently *At*PIP2;1 was also reported to conduct CO_2_ as well as H_2_O and H_2_O_2_ (Wang et al., 2016). Due to the fact that all PIPs have identical selectivity filters (Wallace and Roberts, 2004), which are major determinants of substrate permeability, it is reasonable to assume that other isoforms of the PIP1 and PIP2 subgroups may contribute to CO_2_ diffusion across the plasma membrane and affect *g*_m_ in leaves.

On the molecular scale, the structures and functions of PIPs have been reasonably well described in many plants, but this knowledge is largely limited to the cellular level and scaling it up to the whole plant is more challenging; especially since aquaporin mutants lack an obvious phenotype under standard conditions. Our main aim was to determine the respective roles of three *Arabidopsis* PIP isoforms (*At*PIP2;2, *At*PIP2;4 and *At*PIP2;5) in the regulation of stomatal (*g*_s_) and mesophyll conductance, both of which can substantially limit rates of photosynthesis. At saturating light, the CO_2_ concentration drops to about half that of atmosphere (*c*_a_) at the sites of carboxylation (*c*_c_). The drawdown from *c*_a_ to *c*_i_ (intercellular CO_2_ concentration) is restricted by *g*_s_, accounting for about 60% of the limitation in CO_2_ diffusion, while *g*_m_ accounts for the remaining 40%. Therefore, *g*_m_ is a large, but still poorly understood, limitation of photosynthesis. It has been observed that water deficit has similar effects on *g*_s_ and *g*_m_ (Warren, 2008, 2008), indicating that they are interconnected and, at least in part, controlled by the same mechanisms. Furthermore, since most of the resistance to the diffusion of CO_2_ within the leaf comes from the liquid phase (Warren, 2008), it is conceivable that *g*_m_ could be regulated through the activity of aquaporins. Thus, PIPs may be instrumental in modulating the link between *g*_s_ and *g*_m_. Ultimately, modulating PIP function to increase *g*_m_ without altering *g*_s_, would enhance *A*_net_ without an accompanying increase in transpiration rates and improve the plant’s water use efficiency.

Past studies of the functions of aquaporins have generated a wealth of information concerning single isoforms in *Arabidopsis*, maize and various other herbaceous and woody plant species (Fetter et al., 2004; Zelazny et al., 2007; Ben Baaziz et al., 2012; Gambetta et al., 2013; Li et al., 2015). However, the multitude of different species and methods employed also makes it difficult to develop a cohesive picture of the roles of aquaporins in plants. In this study, we compared three different single knockout mutants of *A. thaliana* and their wild type (WT) to clarify their putative roles in whole-plant water flow. Assigning more clearly defined and specific roles to the different isoforms will aid in determining whether there is redundancy among plant aquaporins. An indication that different aquaporins are not redundant is given by their differing expression patterns. In adult plants, *AtPIP2;2* is highly expressed throughout the plant (Javot et al., 2003; Da Ines, 2010), while *AtPIP2;5* reaches moderate to high levels of expression in mature leaves and guard cells respectively (Alexandersson et al., 2010). However, it shows lower expression levels in roots (Alexandersson et al., 2005), while *AtPIP2;4* is only moderately expressed in leaves but highly expressed in roots (Javot et al., 2003). Since the function of PIPs also depends on their mutual interactions in the tetramer structure (Fetter et al., 2004; Zelazny et al., 2007; Otto et al., 2010), we furthermore examined two double mutants (*pip2;2×2;4* and *pip2;4×2;5*) as well as a triple mutant (*pip2;2×2;4×2;5*).

We grew plants in two environments differing in the ambient relative humidity. This allowed us to compare their responses to different vapour pressure deficits (Vpd), which influence *g*_s_ but not *g*_m_ (Warren, 2008), and distinguish the roles of PIPs in the regulation of *g*_m_ and transpiration. A high Vpd promotes high rates of evaporation and thus triggers plant responses aimed at reducing transpiration such as stomatal closure. Due to the difficulty of observing an obvious phenotype for aquaporin knockout mutants growing under ideal conditions, a high Vpd treatment was applied to increase the relative contribution of aquaporins to plant water flow and the likelihood of finding a distinct phenotypic response.

## Results

### Effects of PIP2 aquaporins on gas exchange under high and low humidity

The difference in evaporative demand between the high humidity (HH) and low humidity (LH) growing conditions had a clear effect on gas exchange and is most strikingly visible in the values of *g*_s_ and its consequences for *A*_net_ (Figure 1A-D). In the LH environment, all plant genotypes displayed 41-61% lower *g*_s_ than in the HH environment (Figure 1C, D). Under LH, however, values of *g*_s_ in the single and double mutant plants lacking *At*PIP2;4 were reduced less than the WT. Under LH, *g*_s_ was significantly reduced in the *pip2;4*, *pip2;2×2;4*, and *pip2;2×2;4×2;5* mutants compared to the WT (*p* = 0.037, 0.049 and 0.008 respectively), whereas no significant differences between the same mutants and the WT were observed under HH conditions. The reduction in *g*_s_ under LH compared to HH was 41%, 53% and 46% for the *pip2;4*, *pip2;2×2;4*, and *pip2;2×2;4×2;5* mutants respectively. The WT *g*_s_ under LH was reduced by 57% compared to HH.

**Figure 1:**
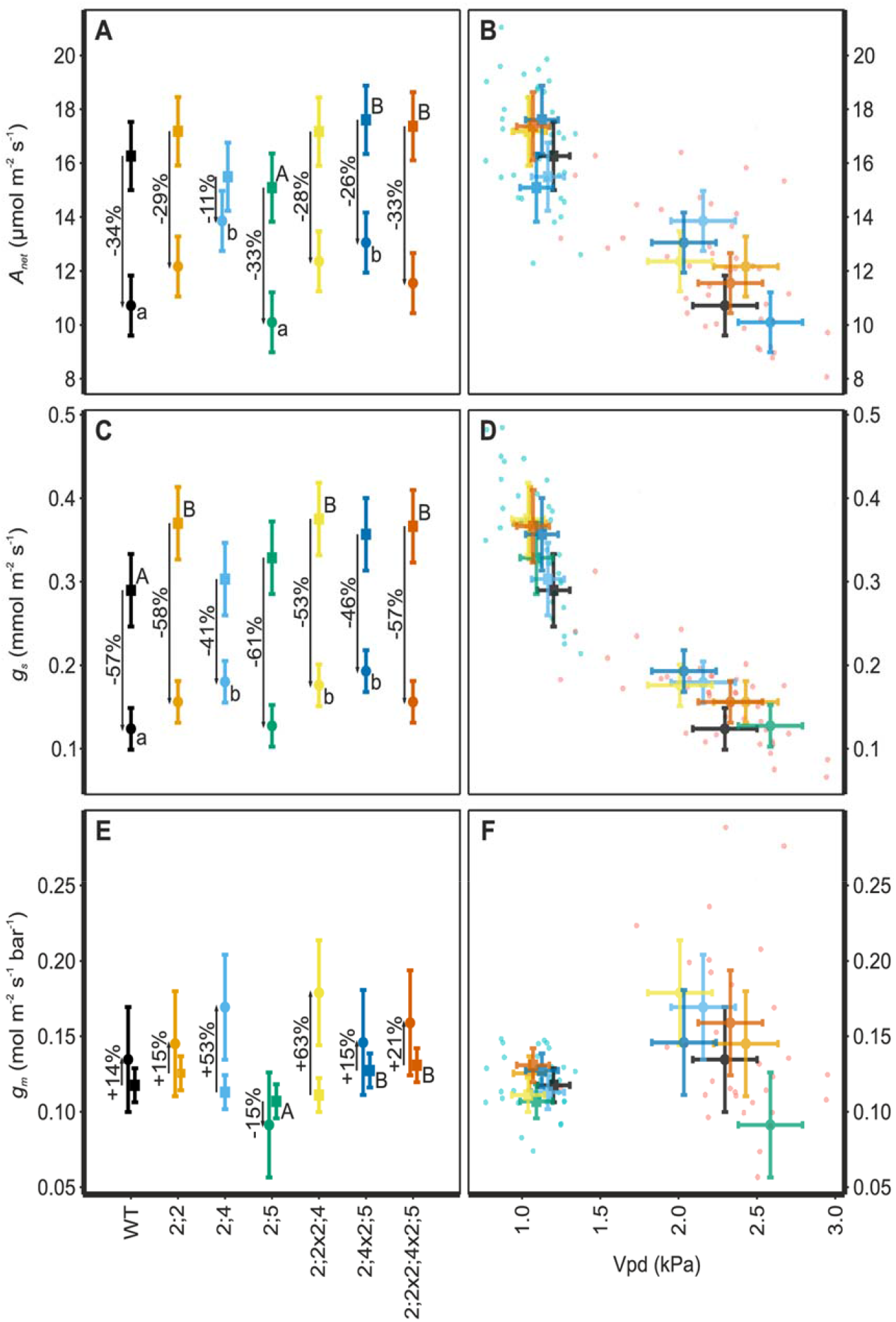
Comparison of gas exchange under low humidity, LH (•), and high humidity, HH (▪). The left-hand panels compare mean ± pooled SE of *A*_net_ (A), *g*_s_ (C) and *g*_m_ (E) among genotypes. Upper and lower case letters indicate statistically significant differences between means within the high and low humidity treatments respectively. The right-hand scatter plots give individual measurements as well as genotype means for gas exchange with respect to Vpd; *A*_net_ (B), *g*_s_ (D) and *g*_m_ (F). The HH plants are shown in blue and LH plants in red. There were no significant differences in the relationship to Vpd among the genotypes within the HH and LH set. Gas exchange parameters are summarised in Table S1.

*A*-*c*_i_ curves confirmed that *A*_net_ was CO_2_-limited under both HH and LH conditions (Figure 2), and was strongly influenced by *g*_s_ and *g*_m_. Consequently, those genotypes in which Vpd had a large effect on *g*_s_ also displayed large differences in *A*_net_ (Figure 1A-D and Table S2). In the WT, *A*_net_ was reduced by 35% in the LH environment compared to the HH environment, whereas the reduction was only 11% for *pip2;4*.

**Figure 2:**
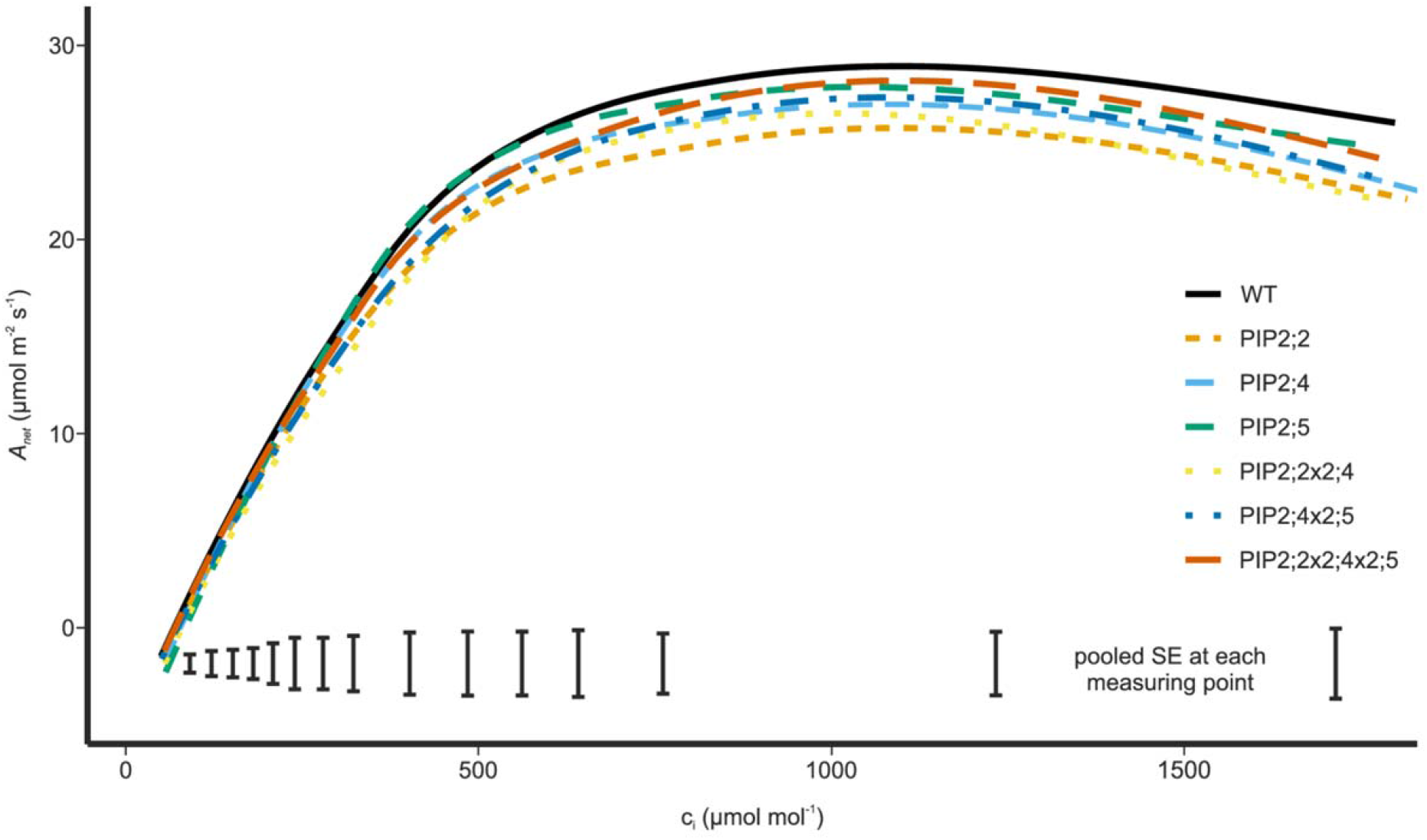
Fitted *A*-*c*_i_ curves for all mutant lines and the WT measured at PAR = 1500 μmol m^−2^ s^−1^. At ambient CO_2_ (≈ 400 μmol mol^−1^ air), *A*_net_ is CO_2_-limited, because this CO_2_ concentration is situated in the linear part of the curve. Shown are fitted means with the pooled SE shown at the bottom of the graph.

In the HH environment, there was a tendency for the genotypes lacking *At*PIP2;2 to have higher values of *g*_s_ in the respective knockout mutants compared to the WT (*pip2;2*, *p* = 0.089; *pip2;2×2;4*, *p* = 0.055; and *pip2;2×2;4×2;5*, *p* = 0.091). Although for the individual mutants alone this was not a statistically significant effect at *p* < 0.05, when all mutant plants lacking *At*PIP2;2 were considered together as a group, the increase of 28% compared to the WT was statistically significant (*p* = 0.034).

Rates of photosynthesis were very uniform among all genotypes under the HH condition (Figure 1A, B). Under LH, however, *A*_net_ declined less in *pip2;4* and *pip2;4×2;5* than in the WT (*p* = 0.010 and 0.041 respectively), as would be expected given their smaller decrease in *g*_s_ from HH to LH compared with the WT. In addition, the *pip2;5* single mutant had lower values of *A*_net_ than the *pip2;4×2;5* double mutant under both growing conditions (LH *p* = 0.028, HH *p* = 0.048).

In terms of *g*_m_, none of the mutant plants differed significantly from the WT under either LH or HH. Noteworthy however, is the fact that unlike *A*_net_ or *g*_s_, *g*_m_ increased slightly under LH in all genotypes except *pip2;5*. Making pairwise comparisons, we found that *pip2;5* had a significantly lower *g*_m_ than *pip2;4×2;5* and *pip2;2×2;4×2;5* under HH (*p* = 0.065 and 0.049 respectively, Figure 1E, F), whereas under LH differences were smaller and not statistically significant (*p* = 0.217 and 0.075 respectively, Figure 1E, F). There was a similar pattern of effects between the HH and LH environments for *A*_net_ among these pairs of mutants: *pip2;5* vs. *pip2;4×2;5* and *pip2;5* vs. *pip2;2×2;4×2;5* (LH *p* = 0.028 and 0.280 respectively, HH *p* = 0.048 and 0.088 respectively).

Both, *A*_net_ and *g*_s_, responded to Vpd and the individual plant measurements as well as the means for each genotype form two distinct clusters (Figure 1B, D, F). Values of *g*_m_ were less responsive to Vpd, but nevertheless form two clearly separate clusters. Variation in terms of Vpd was larger under LH (2.0 −2.5 kPa) than HH (1.0 – 1.25 kPa), but none of the mutants differed significantly from the WT in their Vpd under either growing condition, and thus Vpd does not account for statistically significant differences observed in *A*_net_, *g*_s_ or *g*_m_.

### Effects of PIP2 aquaporins on mesophyll conductance under high and low humidity measured through A-response curves

We measured *A*-*c*_i_ curves (Figure 2) to detect effects of the PIP2 mutations on the limits of CO_2_ fixation. There were no significant differences among the genotypes for either the maximum rate of electron transport (*J*_max_) or the maximum rate of carboxylation (*V*_cmax_) (Table S2). Furthermore, there were no differences in *c*_i_/*c*_a_ at low CO_2_ (< 400 μmol CO_2_ mol^−1^ air), but at high CO_2_ (> 400 μmol CO_2_ mol^−1^ air), *c*_i_/*c*_a_ was significantly lower in all mutants lacking *At*PIP2;5 than in the WT (*p* = 0.008, 0.013 and <0.001 for *pip2;5*, *pip2;4×2;5* and *pip2;2×2;4×2;5* respectively; Figure 3A). The *A*-*c*_i_ curves of the remaining mutants (*pip2;2*, *pip2;4* and their double mutant *pip2;2×2;4*) were determined to be significantly different in shape from the WT using a mixed effects generalised additive model (GAMM, *p* = 0.002, 0.039 and 0.042 respectively), but this difference was not large enough to affect *J*_max_ or *V*_cmax_.

**Figure 3:**
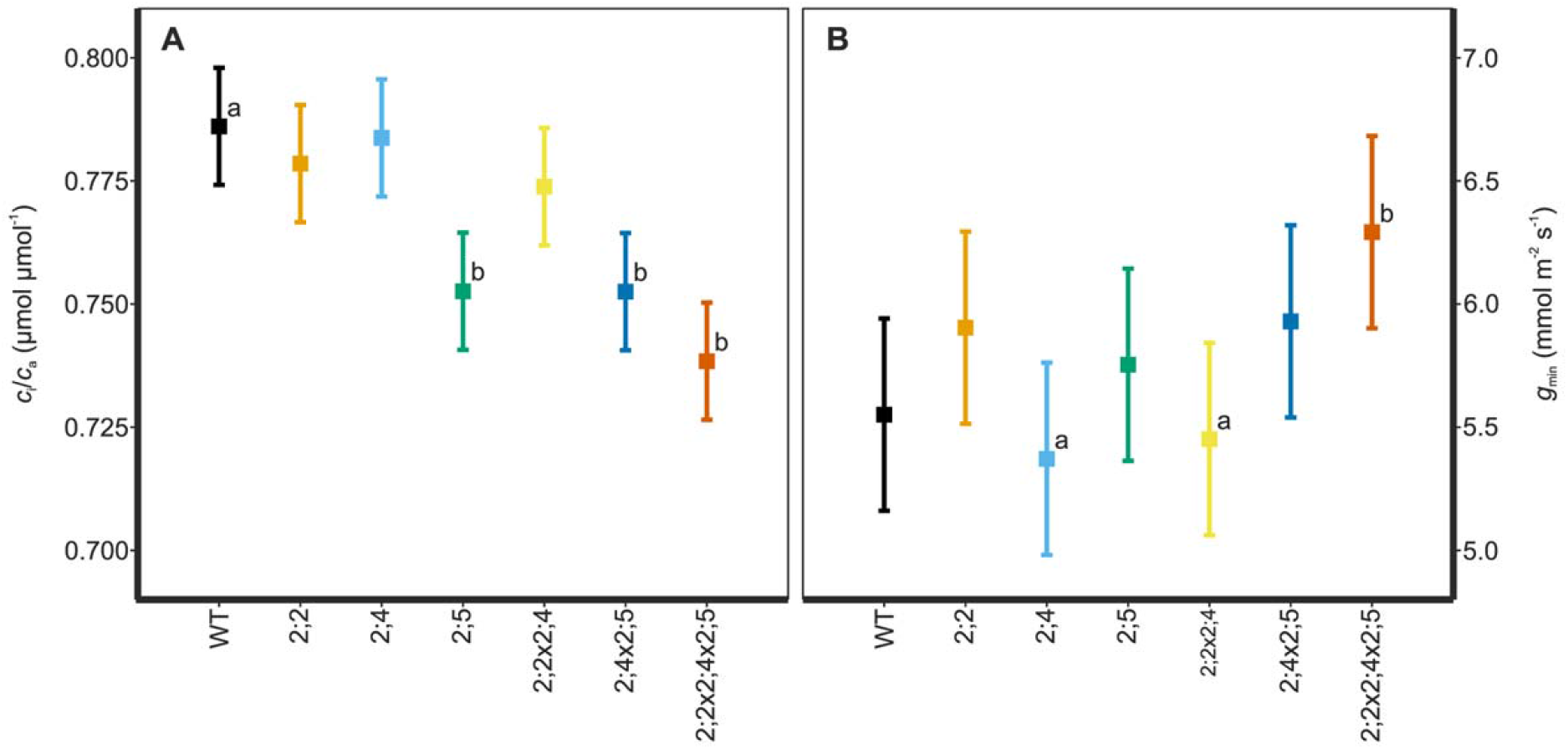
A – The mean ± pooled SE *c*_i_/*c*_a_ for measurement points of the *A*-*c*_i_ curves at higher than ambient CO_2_ concentrations (≥ 400 μmol mol^−1^ air) and under high humidity. Letters indicate statistically significant differences compared to the WT. B – The mean ± pooled SE for minimum conductance, *g*_min_, under HH. Letters indicate statistically significant differences between mutants.

Figure 4 shows the light response curves for LH-grown plants measured under low oxygen. In line with the steady-state measurements of gas exchange, *pip2;5* produced light response curves with lower values of *A*_net_, as well as *g*_s_, than the other genotypes (13% and 5% lower respectively compared to the WT). At points on the curve at saturating PAR (500-2000 μmol m^−2^ s^−1^ for these plants), *A*_net_ was not significantly different from the WT for any of the genotypes. However, for measurement points at sub-saturating PAR (< 500 μmol m^−2^ s^−1^), *A*_net_ of *pip2;5* was lower than that of all other genotypes including the WT (21 – 50% lower, *p* ≤ 0.018). At low light intensities, *g*_s_ did not differ among genotypes, whereas at higher light intensities (1000-2000 μmol m^−2^ s^−1^), *pip2;5* differed significantly from all other genotypes including the WT. Using GAMM, we were able to confirm these differences in shape of the light-response curves, which were significant for *A*_net_ as well as *g*_s_ in *pip2;5* compared to the WT (*p* = 0.043 and 0.001 respectively). In addition, the curves *pip2;2* and *pip2;2×2;4×2;5* also differed significantly from the WT in their response of *g*_s_ to increasing PAR (*p* = 0.049 and 0.027 respectively).

**Figure 4:**
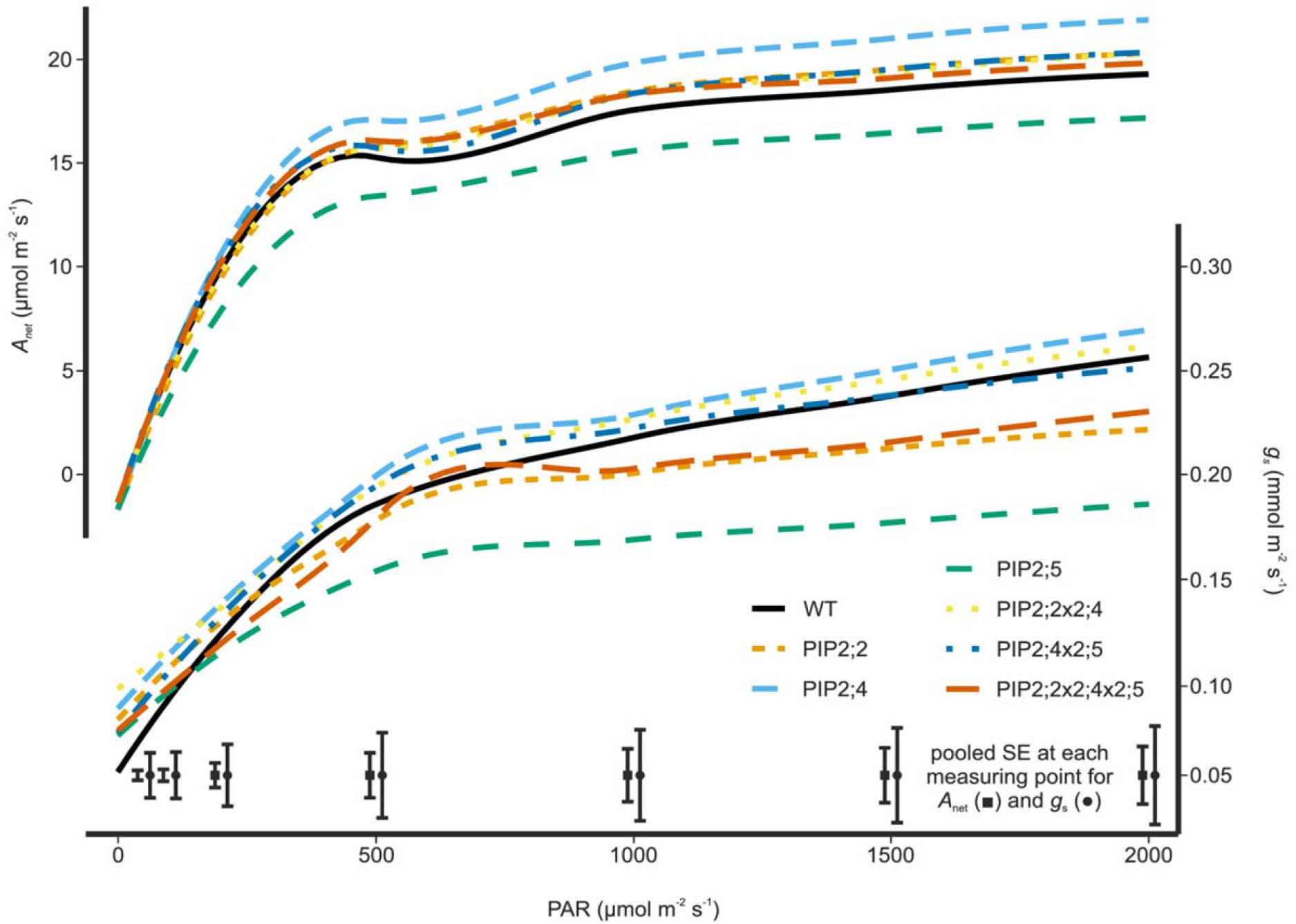
Light response curves measured in under LH showing the rate of photosynthesis (above) and the stomatal conductance (below) in response to increasing radiation. In both cases the *pip2;5* mutant stands out displaying 13% lower *A*_net_ as well as a much slower and 5% smaller response of *g*_s_ compared to the wild type. *Pip2;4* showed the opposite trend with 21% higher *A*_net_ and 19% higher *g*_s_ compared to the wild type. Given are fitted means with the pooled SE at the bottom of the graph.

Values of the minimum conductance of water (*g*_min_) (Figure 3B) were within the expected range of 5-10 mmol m^−2^ s^−1^ which is small compared to the *g*_s_ of fully open stomata. No significant differences in *g*_min_ were found between the WT and any of the mutants. Only *g*_min_ of *pip2;2×2;4×2;5* tended to be higher than the WT (*p* = 0.065), and furthermore both *pip2;4* and *pip2;2×2;4* differed significantly from *pip2;2×2;4×2;5* (*p* = 0.029 and 0.037 respectively).

### CO_2_ conductance of AtPIP2;5 expressed in yeast

Yeast cells expressing either *At*PIP2;5, *At*CA1 or both and loaded with fluorescein diacetate displayed significantly different fluorescence intensities after the application of the CO_2_ mixing buffer (Figure 5). Entry of CO_2_ into yeast cells expressing *At*CA1 resulted in H_2_CO_3_^−^ formation (and subsequent dissociation into H^+^ and CO_3_^−^) and thus intracellular acidification as indicated by a decrease in fluorescence intensity. Fluorescence intensity did not decrease in yeast cells expressing only *At*PIP2;5, because *At*CA1 was not present and thus no significant acidification occurred (Figure 6). This assay was used as the first control. Yeast cells expressing only *At*CA1 were used as a second control in order to quantify the CO_2_ permeability of the yeast membrane in the absence of *At*PIP2;5. These cells displayed a slight decrease in fluorescence intensity due to CA1-facilitated formation of H_2_CO_3_^−^ and subsequent intracellular acidification (Figure 6). When *At*PIP2;5 and *At*CA1 were co-expressed, CO_2_ entry into the cells was >100-fold more rapid than in the controls (Figure 6) indicating that the CO_2_-permeability of the membrane was drastically increased by the insertion of *At*PIP2;5.

**Figure 5:**
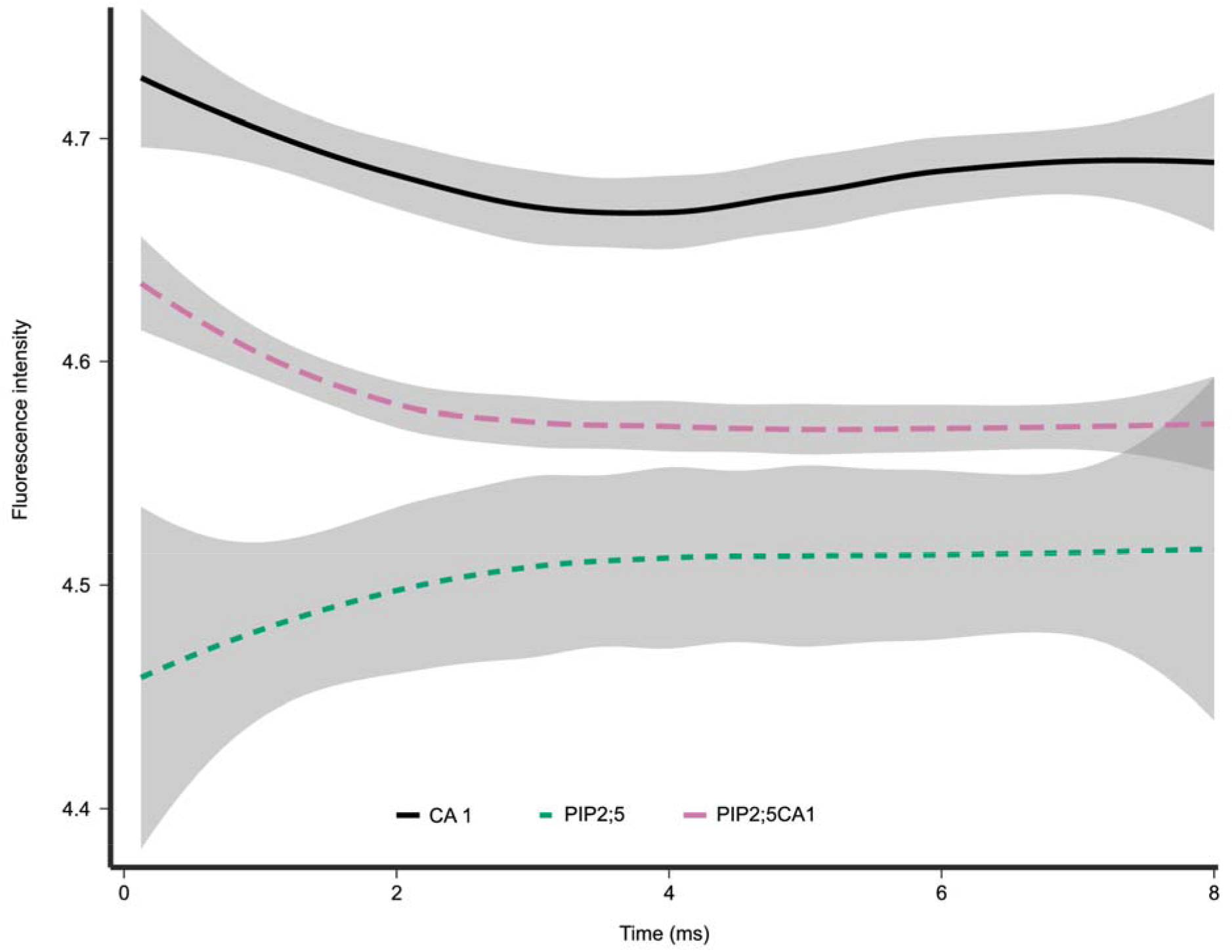
Fluorescence intensity for yeast cells loaded with fluorescein diacetate measured at 0.125 ms intervals. Intracellular acidification in response to the entry of CO_2_ causes a decrease in the fluorescence intensity of yeast cells. Given are average curves with 95% confidence intervals.

**Figure 6:**
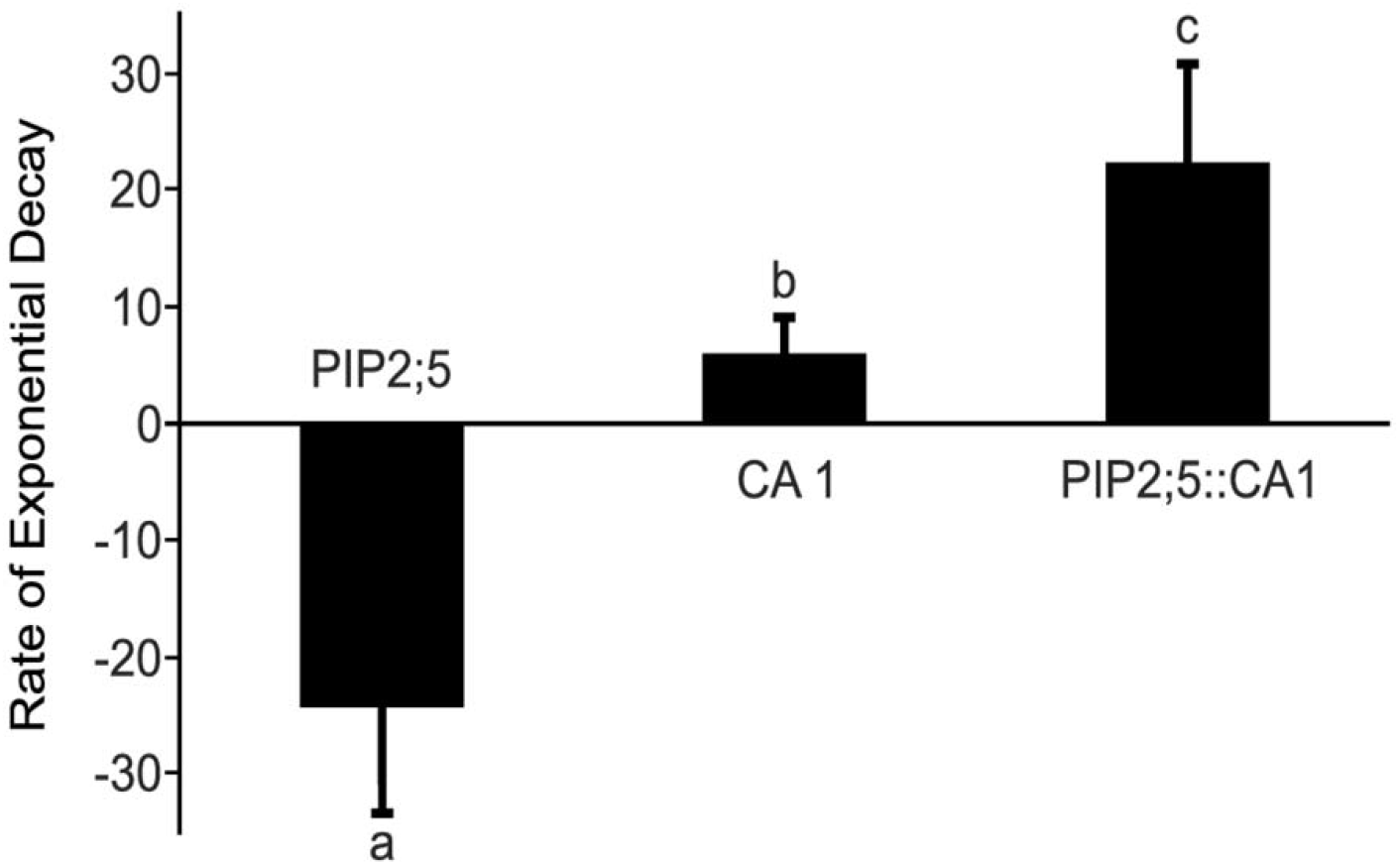
CO_2_-induced intracellular acidification rate of *S. cerevisiae* cells expressing *At*PIP2;5, *At*CA1 or both. Yeast cells were exposed in a ratio of 1: 1 (v/v) to a CO_2_-mixing buffer (25mM HEPES, 75mM NaHCO3, pH 6). Kinetics of acidification were measured with an excitation wavelength of 460 nm and emission above 515 nm using a stopped-flow spectrophotometer. Bars represent the CO_2_ permeability of yeast expressed as exponential decay rate of fluorescence intensity. The kinetics of the decrease in fluorescence were obtained by fitting an exponential decay function to the curves shown in Figure 5 in order to calculate the rate constants. Values are means ± SD of 3 replications. Different letters denote statistically different values at *p* < 0.05.

## Discussion

In the *Arabidopsis* plants grown with an ample water supply and under HH conditions, similarly to earlier reports (Javot et al., 2003; Wang et al., 2016), water flow through the various tissues did not appear to be affected by PIP knockout mutations as evidenced by the very uniform values of *A*_net_ and *g*_s_. Increasing the evaporative demand by lowering ambient air humidity causes an increase in the water flow through the plant due to a higher Vpd and hence the contribution of aquaporins to the regulation of water flow through the plant also increases. By allowing plants to develop and grow under LH conditions, we aimed to amplify any differences in water relations between the WT and knockout mutants (Figure 1).

*AtPIP2;5* has been reported to respond to various stresses such as H_2_O_2_ (Hooijmaijers et al., 2012) and salt (Lee and Zwiazek, 2015), as well as being up-regulated by drought (Alexandersson et al., 2010), but it is only expressed at intermediate levels under standard growing conditions, especially in the roots (Jang et al., 2004; Lee et al., 2012). This is consistent with the absence of an effect on *A*_net_ and *g*_s_ in *pip2;5* compared with the WT in our HH environment. However under LH, *g*_m_ was reduced at steady state (Figure 1E) as was *A*_net_ and *g*_s_ in the light-response curves (Figure 4) in this mutant compared to the WT. Reduced *g*_m_ is a commonly reported response to drought (Warren et al., 2004; Warren, 2008, 2008), but since our plants were not experiencing soil water stress, *g*_m_ remained unchanged or slightly increased under LH compared with HH. We suggest that high soil water content in our experiment allowed plants to avoid water-stress under LH partially explaining the lack of a concomitant effect on *A*_net_ or *g*_s_ in *pip2;5* with reduced *g*_m_. The differences in *g*_m_ that we found suggests that *At*PIP2;5 is involved in the regulation of *g*_m_ *in planta*. Our stopped-flow measurements on yeast cells expressing *At*PIP2;5 provided supporting evidence for this, since they clearly show that *At*PIP2;5 is permeable to CO_2_, when expressed in yeast, and thus its ability to regulate *g*_m_ is likely due to it directly facilitating CO_2_ diffusion across the cell membrane. *At*PIP2;5 has not previously been shown to alter CO_2_ fluxes across cell membranes to affect *g*_m_ nor has it, to our knowledge, been tested for CO_2_ permeability.

The fact that *AtPIP2;5* is upregulated during drought, differentiates it from most other PIPs and the lack of functional *AtPIP2;5* in combination with LH may thus have triggered the drop in *g*_m_. It is furthermore only weakly co-expressed with one other PIP isoform, *At*PIP1;4 (Alexandersson et al., 2010), which reduces the likelihood of other aquaporins compensating for its lack in the knockout mutants, but in the very same mutants, the proper insertion of *At*PIP1;4 into the plasma membrane may be reduced due to the missing interaction with *At*PIP2;5. *At*PIP2;5 may thus also be involved in maintaining *g*_m_ across a greater range of watering and humidity regimes by affecting the function of *At*PIP1;4 (Fetter et al., 2004; Zelazny et al., 2007; Otto et al., 2010), which is not only upregulated by drought (Alexandersson et al., 2010), but has been shown to also contribute to CO_2_ membrane permeability (Li et al., 2015). Further evidence to support this theory is provided by the results of our light-response curves where *g*_s_ was similar for all genotypes, whereas the *A*_net_, of *pip2;5* was clearly lower than in the other genotypes. Therefore, contrary to our expectations, *g*_s_ did not behave like *A*_net_ in *pip2;5* in response to a changing light environment, particularly at sub-saturating PAR. It would therefore appear that *At*PIP2;5, despite its relatively low abundance (Lee et al., 2012), does reduce the resistance to CO_2_ diffusion through the mesophyll under light- and CO_2_-limiting conditions.

Further evidence in support of *At*PIP2;5 regulating *g*_m_ is provided by our *A*-*c*_i_ curves. Its knockout mutation did not appear to significantly affect *J*_max_ or *V*_cmax_, which was not unexpected since none of the PIPs have previously been shown to impact chlorophyll fluorescence or CO_2_ fixation by Rubisco. However, the *c*_i_/*c*_a_ was significantly lower than the WT at high [CO_2_] for all mutants lacking *At*PIP2;5 (Figure 3A). Since *c*_a_ was constant, the lower *c*_i_/*c*_a_ was due to lower *c*_i_, likely the result of an incremental effect on a combination of *A*_net_ and *g*_s_ during the *A*-*c*_i_ curve or an increase in *g*_m_.

The interaction of PIPs was previously demonstrated to modify their function and export from the ER (Fetter et al., 2004; Zelazny et al., 2007; Otto et al., 2010). PIP2s interact with PIP1s forming tetrameric assemblies in the ER to ensure proper export and insertion into the plasma membrane. PIP1s are retained in the ER if no interaction with other PIP isoforms occurs (Zelazny et al., 2007) and it is thus conceivable that *At*PIP2;5, even though normally weakly expressed, impacts membrane permeability to CO_2_ by providing targeting signals for the proper integration of CO_2_-conducting PIP1s, *At*PIP1;4 in particular, into the plasma membrane. The fact that *pip2;5* displayed significantly lower values of *A*_net_ and *g*_s_ than the respective double and triple mutant furthermore suggests that its role as a targeting signal for proper membrane integration of PIP1s can be fulfilled by other PIP2s that are activated/upregulated in multiple knockout mutants. Serving as targeting signals is thus most likely a function common to all PIP2 isoforms. In general, multiple knockout mutations of PIP2s did not cause larger effects than single knockout mutation, indicating that the remaining intact isoforms are capable of compensating under adequate watering conditions.

The *g*_s_ of *pip2;2*, *pip2;4* as well as that of the double and triple mutants increased relative to the WT. Significant differences occurred for *pip2;2*, *pip2;2×2;4* and *pip2;2×2;4×2;5* under HH, whereas for *pip2;4*, *pip2;2×2;4* and *pip2;4×2;5* higher *g*_s_ was observed under LH. *At*PIP2;2 has been more intensely studied than *At*PIP2;4 or *At*PIP2;5, and has been found to be among the most abundantly expressed PIPs in the plant (Javot et al., 2003; Da Ines, 2010). Its knockout mutation has furthermore been reported to induce no compensatory aquaporin gene expression nor a visible phenotype (Javot et al., 2003). However, *At*PIP2;2 contributes to cell hydraulic conductivity in the root cortex (Javot et al., 2003) and thus, without the up-regulation of other PIPs or a visible phenotype, it is likely that any compensatory response occurs at the level of stomatal regulation leading to the higher *g*_s_ that we report. This in turn would enhance the water flow through plant tissues via other PIP isoforms or the apoplast as may also be the case in the *pip2;4* mutant.

Interestingly, under LH *g*_s_ of mutants lacking *At*PIP2;2 did not significantly differ from the WT. *AtPIP2;2* has previously been shown to be drought-sensitive (Alexandersson et al., 2010), but as our plants were not drought stressed, it is probable that the absence of *At*PIP2;2 may be compensated by increased water flow through the apoplastic pathway or other PIPs. *At*PIP2;4 appears to be a better candidate for manipulating plant water relations, since *pip2;4* displayed higher *g*_s_ compared with the WT in the LH environment, but not under HH.

None of the plants that we investigated showed differences in *g*_min_ (Figure 3B) or RWC (Table S2), which indicates that the observed effects on *g*_s_ and *g*_m_ are unlikely to be due to an effect of the knockout mutation on leaf water status. We also did not find any visible phenotype under either HH or LH. Therefore, the differences we report between the mutants and the WT are more likely to be a direct result of the knockout mutation and lack of aquaporin function rather than an indirect effect caused by altered leaf water status.

## Conclusion

We report that *At*PIP2;5 is permeable to CO_2_ when expressed in yeast and that it is a physiologically relevant regulator of mesophyll conductance of CO_2_ in leaves of *A. thaliana* under conditions of high evaporative demand.

Likewise, *At*PIP2;4 plays a role in maintaining a positive leaf water balance and maintaining high IWUE. It may also be a suitable target for crop improvement, since the lack of *At*PIP2;4 caused a 29% increase in *A*_net_ when Vpd was high, though at the expense of IWUE. Identification of the mechanisms underpinning this result may represent the means of teasing apart the factors regulating *g*_s_ and *g*_m_.

## Materials and Methods

### Plant Material

T-DNA single knock-out mutants of *Arabidopsis thaliana* were obtained from the Nottingham Arabidopsis Stock Centre (NASC – www.arabidopsis.org) for the following aquaporins: *PIP2;2* (N871747), *PIP2;4* (N105980) and *PIP2;5* (N117303) in the Columbia background (Alonso et al., 2003). All genotypes were checked by PCR to confirm correct T-DNA insertion and homozygosity. Only homozygous plants were used to grow a seed stock and in the subsequent experiments. Multiple knock-out plants were created by crossing, which resulted in a total of two different double mutant lines – *pip2;2×2;4*, *pip2;4×2;5* – and one triple mutant – *pip2;2×2;4×2;5*.

Since *g*_m_ exhibits acclimation to environmental conditions (Warren, 2008), we grew the plants under different humidities in two locations instead of raising them in the same environment and subjecting them to short-term treatments, which would likely affect both *g*_m_ and *g*_s_ and thus prevent us from separating the role of aquaporins in these two processes. The environmental parameters for the high humidity and low humidity locations are summarised in Table S1.

### Low humidity growing conditions

The experiment was carried out in a greenhouse at the University of Sydney, Sydney, Australia. Seeds of the *A. thaliana* genotypes were sown in pots containing a potting mix (Scotts Osmocote – plus trace elements) and germinated in the light under the conditions described in Table S1. Seedlings were transplanted four days after germination into square plastic pots 20 cm high and 6 cm wide. Pots were over-filled with the same pre-fertilised potting mix, as described in Flexas et al. 2007 (Flexas et al., 2007). Young seedlings were kept under a transparent plastic cover for a few days after transplanting to help keep them moist. The pots were arranged randomly and rearranged every second day to ensure even light interception and minimise any effects of air movement caused by the air conditioning. All measurements were carried out between 9:30 am and 16:30 pm on 25-39-day old plants using only fully expanded leaves at least 4 cm long, a period when *A*_net_ is reported to be stable (Flexas et al., 2007). The experiment was conducted during the Australian spring from mid-October and mid-December 2015. During this time, new seeds were planted at weekly intervals in order to continuously provide plants of equivalent ages for measuring.

Plants were provided with ample water in order to prevent soil water stress signalling from the roots, because our aim was to expose plants to low air humidity without imposing other accompanying stresses. Water was provided from below as soon as the soil surface had dried (every 2-3 days), but after the day’s measurements.

### High humidity growing conditions

Seeds of the *A. thaliana* genotypes were sown and grown in a pre-fertilised peat-vermiculite mixture (1:1) in square 8 cm wide and 6 cm high plastic pots. Seedlings were transplanted four days after germination into over-filled pots as described above. Plants were grown in a growth chamber under conditions described in Table S1 and the light spectrum shown in Figure S1. Measurements were carried out between 9:00 am and 16:00 pm from October 2016 until January 2017 on 27-33-day old plants using only fully expanded leaves at least 4 cm long. The plants were watered as in the low humidity treatment described above, but due to the higher air humidity in these growing conditions, watering was required only once a week.

### Gas Exchange Measurements

All gas exchange measurements were carried out with the portable photosynthesis system LI6400XT infra-red gas analyser (IGRA) equipped with a fluorescence chamber (LI-COR Biosciences, Nebraska, USA) during the same 7-h time-window every day in order to minimise diurnal effects. The leaf chamber (leaf chamber fluorometer – LCF), a 2 cm^2^ circular cuvette, allowed single leaves to be measured. A few leaves that did not fill the chamber entirely, were placed in the middle of the circular window leaving equal gaps on either side. The area of these gaps was calculated using the following formula:

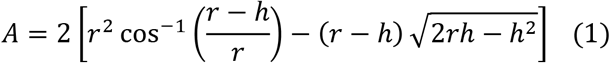
where *r* is the chamber radius (7.979 mm) and *h* is the height of the circle segment not covered by the leaf or in other words, the distance from the edge of the leaf to the rim of the chamber. *h* was obtained by:

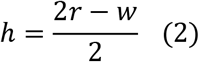
where *w* is the width of the flat leaf as measured with a sliding calliper. The area of the chamber covered by the leaf was then calculated simply subtracting the area of the two gaps as calculated above from the total area of the leaf chamber (2 cm^2^).

In the low humidity greenhouse experiment, photosynthetic light responses were measured under non-photorespiratory conditions (1% O_2_, 400μmol mol^−1^ CO_2_) to determine the relationship between photosynthetic rate and light intensity as well as the relationship between chlorophyll fluorescence (*J*_f_) and the rate of electron transport (*J*_a_). High purity N_2_ gas (BOC Gas, Australia) was mixed with air to create a 1% O_2_ mixture, directly attached to the air inlet of the LICOR-6400. Under non-photorespiratory conditions, *J*_a_ is entirely dependent on gross photosynthesis *A*:

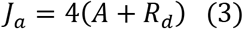

Before each measurement, the plant leaf was dark adapted for 30 min by wrapping it in aluminium foil. The curves were subsequently measured using an automated program with following fixed settings: T: 25°C, leaf fan: fast, CO2R: 400 μmol mol^−1^, flow: 300 μmol s^−1^ and 10% blue light. At the beginning of the program the leaf was given a further 5 min of dark adaptation within the chamber. This step allowed the measuring chamber-leaf system sufficient time to reach a steady state. Light adaptation lasted for 30 min at maximum irradiance (PAR 2000 μmol m^−2^ s^−1^). The light curves began at the highest irradiance and decreased at ca. 4 min intervals of acclimation time between each step: 2000, 1500, 1000, 500, 200, 100, 50, 20 and 0 μmol m^−2^ s^−1^.

In the high humidity growth chamber experiment, full *A*-*c*_i_ curves were measured with an automated program throughout the curve with the following fixed parameters: flow: 300 μmol s^−1^, T: 25°C, PAR: 1500 μmol m^−2^ s^−1^, 10% blue, leaf fan: fast, CO_2_R: 400 μmol mol^−1^. Plants were allowed an acclimation time of 10 min in the leaf chamber before the first measurement at 400 μmol mol^−1^ CO_2_. After the first measurement, the following steps with 3 min acclimation time between each step were used to obtain a complete *A*-*c*_i_ curve: 450, 550, 650, 750, 850, 1000, 1500, 2000, 400, 350, 300, 250, 200, 150, 100, 50 μmol mol^−1^ (Warren and Dreyer, 2006).

The Laisk method (Laisk, 1977; Brooks and Farquhar, 1985) was used to estimate *c*_i_* (photorespiratory compensation point) and R_d_ (respiration in the light), which are variables required for the calculation of *g*_m_. In both experiments (LH and HH conditions), an automated program with the same general settings was used for the Laisk method: T: 25°C, leaf fan: fast, CO2R: 400 μmol mol^−1^, flow: 200 μmol s^−1^, PAR 1500 μmol m^−2^ s^−1^, 10 % blue. Before the measurements for the Laisk method, plants were allowed to acclimate and then measured at steady state. The *A*-*c*_i_ curves were measured at PAR 300, 150, 100, 50 μmol m^−2^ s^−1^ and after each curve, CO_2_ was returned to 400 μmol mol^−1^ for 5 min to maintain RuBisCo activation. The CO_2_ steps used for the curve are 150, 125, 100, 75, 50 μmol mol^−1^ with ca. 3 min acclimation time between each step.

Water loss through leaves is not only governed by stomata, but also occurs through the epidermis and cuticle. *g*_min_ describes the minimum conductance that is the rate of water loss from leaves due to direct diffusion through the cuticle and leaky stomata. This pathway for gas exchange is often neglected as it represents only values in the range of 5-10 mmol m^−2^ s^−1^ H_2_O (Figure 3B, Table S2, (Duursma et al., 2019)) as compared to a *g*_s_ of 100 - 400 mmol m^−2^ s^−1^ for actively transpiring *Arabidopsis* leaves (Figure 1C). Nevertheless, neglecting *g*_min_ introduces an error in estimations of *g*_s_. Aquaporins may have the potential to alter *g*_min_ either through the regulation of leaf development or by increasing/decreasing the water permeability of the epidermis itself. Therefore, all gas exchange data were adjusted to account for *g*_min_ as well as any CO_2_ leaks into or out of the LICOR measuring chamber. We estimated *g*_min_ for fully expanded leaves using the protocol described by Sack (Sack, 2010) with some modifications to accommodate fragile and small *Arabidopsis* leaves: for each data point, three leaves that were suitably large for gas exchange measurements were excised from 21 plants/line close to the centre of the rosette. The leaves were weighed for initial fresh weight (*FW*), placed on millimetre graph paper and photographed to calculate their initial leaf area using ImageJ as described by Wang (2016) (Wang, 2016). After photographing, they were placed on labelled Petri dishes and allowed to dry at room temperature (20°C) for 1 h until complete stomatal closure. From this point on, leaves were weighed every 25-30 min and at each weighing step, temperature, ambient air humidity and time was recorded. The first time point after the 1 h drying period was taken to be time 0. After six to ten time points had been obtained over the course of 3-4 h, the leaves were again photographed to calculate their final leaf area. The collected data were input into the “g_min_ Analysis Spreadsheet Tool” (Sack, 2010) together with the saturation vapour pressure (Table A7, p. 430-431, Pearcy et al. 1989 (Pearcy R.W., 1989)) in order to calculate *g*_min_.

The leak flow was calculated using the manufacturer’s instruction, however, we placed an intact leaf in the chamber instead of carrying out the estimation for an empty chamber. In the dark, the leaf’s respiration rates should not be affected by changing CO_2_ concentrations or flow rates and thus can be considered constant. Therefore, we were able to obtain a diffusion coefficient of 0.40 mol s^−1^ for the actual measuring conditions, which is similar to the 0.44 mol s^−1^ obtained by Flexas et al. (2007) (Flexas et al., 2007).

The mesophyll conductance of CO_2_, was estimated using the variable J method as described by Harley et al. 1992 (Harley et al., 1992):

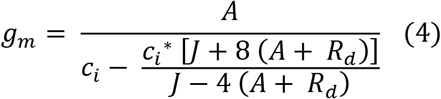

The variables *A*, *c*_i_ and *J* were obtained from the combined gas exchange and chlorophyll fluorescence measurements. R_d_ as well as *c*_i_* were estimated from the Laisk method by plotting the *A*-*c*_i_ curves in excel and calculating their common intersection point using slope-intercept regression (Walker and Ort, 2015). As we did not find any statistically significant differences in either R_d_ or *c*_i_* between the different genotypes, we calculated a global average for both variables for each growing condition in order to obtain a more robust estimate of *g*_m_.

Parameters *J*_max_, *V*_cmax_ and the inflection point were estimated from *A*-*c*_i_ curves. These parameters were obtained using the online Excel tool provided by Carl Bernacchi’s lab (Bernacchi), which is based on the Michaelis-Menten constants of CO_2_ (*K*_c_) and O_2_ (*K*_o_) at *c*_i_ as described by Bernacchi et al. (2001) (Bernacchi et al., 2001).

### Leaf water relations

The same harvested leaves that had been used for estimating cuticular conductance were floated on water overnight to obtain their saturated weights (*SW*). Finally, the leaves were dried in a drying oven overnight at 60°C to obtain their dry weights (*DW*). The relative water content (*RWC*) was then calculated using the following formula:

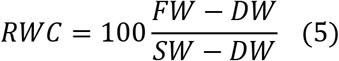

### RNA Extraction and cloning

Tissue from 6 days old seedlings of *A. thaliana* ecotype Columbia (Col-0) were used for RNA extraction. Total RNA was extracted using the QIAGEN RNeasy Plant Mini Kit (Qiagen, Valencia, CA, USA). RNA concentration and purity were assessed using the Thermo Scientific™ NanoDrop™ One Microvolume UV-Vis Spectrophotometer (Thermo Scientific, Waltham, MA, USA). The first-strand cDNAs were synthesized from 1 μg of total RNA using Superscript II reverse transcriptase (Invitrogen) and an oligo (dT) primer. Coding sequences of *Arabidopsis* PIP2;5 (AT3G54820) and carbonic anhydrase 1 (CA1; AT3G01500) were amplified with Phusion DNA Polymerase using the primers listed in Table S3. The PCR products of the expected size were eluted from the gel and purified using the Wizard® SV Gel and PCR Clean-Up System (Promega, Madison, WI, USA). The PCR products were then cloned into a pCR2.1-TOPO vector using the Topo TA Cloning kit (Invitrogen, Carlsbad, CA, USA) and transformed into DH5α chemically competent cells (Invitrogen, Carlsbad, CA, USA). About three to six white colonies were sequenced for each PCR product using M13 sequencing primers.

### Plasmid construction and yeast transformation

Open reading frame (ORF) of *At*PIP2;5 and Carbonic anhydrase 1 (*At*CA1) were cloned as Gateway entry clones in plasmid pDONR221 (Invitrogen). ORFs of *At*PIP2;5 and CA1 from the entry clones were shuttled into the galactose-inducible yeast expression plasmid pAG426GAL-ccdB (Addgene: Plasmid #1415) and pAG425GAL-ccdB (Addgene: Plasmid #14153), respectively by Gateway LR cloning reaction. The S.c. EasyComp™ Transformation Kit (ThermoFisher Scientific) was used to transform *S. cerevisiae* yeast strain (INVSc1 from ThermoFisher Scientific) with each plasmid DNA. Double transformants were used for CO_2_ permeability measurements containing *At*CA1 and *At*PIP2;5 (AtCA1::AtPIP2;5) constructs were selected by ura3 and leu2 complementation. Expression of the constructs was verified by qRT-PCR using SYBR Green I dye in an Applied Biosystems 7500 Fast system.

### Loading of yeast cells with fluorescein diacetate

Loading of the yeast cells with fluorescein diacetate was carried out according to the protocol described by Bertl and Kaldenhoff (2007) (Bertl and Kaldenhoff, 2007). In brief, cells were harvested by centrifugation and then resuspended in loading buffer (50 mM HEPES-NaOH pH 7.0; 5 mM 2-deoxy-D-glucose) with 50 μM of fluorescein diacetate, incubated for 14 min at 30 °C with shaking at ~ 225 rpm and centrifuged again at 1,700 g for 3 min at 4 ° C. The cells were then resuspended in incubation buffer (25 mM HEPES; 75 mM NaCl) and kept on ice until use.

### CO_2_ conductance measurements

The entry of CO_2_ through the plasma membrane was followed by intracellular acidification and decreased fluorescence in whole yeast cells loaded with fluorescein diacetate. The cells were resuspended in incubation buffer to a final OD600 of 60 just before use. 50 μL of dissolved cells in the incubation buffer were mixed rapidly in a 1: 1 (v / v) ratio with CO_2_-Mixing Buffer (25mM HEPES, 75mM NaHCO3, pH 6) at a rate of 100 μL / s in a stopped flow spectrophotometer (Applied Photophysics, DX.17 MV). The CO_2_ input was followed by the decrease in fluorescence intensity (90º). The spectrophotometer emitted at a λ of 490 nm (maximum excitation wavelength of the fluorescein). The receiver had a filter attached that did not allow the passage of wavelengths below 515 nm, because fluorescein emits at λ of no longer than 514 nm. The fluorescence was recorded over time and the conductivity quantification (K-relative) was calculated by fitting the experimental data to a function of decreasing exponential during the first 8.0 ms using SigmaPlot 11.0 (Systat Software Inc., Chicago, IL, USA).

### Data processing and statistical analysis

ANOVAs were conducted separately for the LH and HH experiment in R (package Deducer) using a linear model with plant genotype and the measured variable as the factors, and for each graph/panel we calculated the pooled standard error:

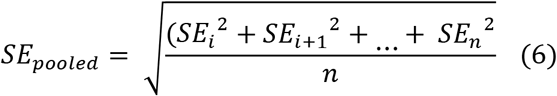
where SE_*i*_ is the standard error of the mean and *n* is the total number of means. *n* = 21 for estimations of *g*_min_ and RWC, while for all gas exchange measurements *n* = 4 - 9. Tukey’s multiple comparison was used to compare the means of all measured variables for the mutant lines to the WT as well as with each other.

GAMM (package = ‘mgcv’) (Wood, 2017b) was used to evaluate the photosynthetic response curves with the mutant line as a parametric term and a smoothing term for PAR and *c*_i_. Due to heterogeneous variation we employed the weighting function ‘weights=varExp’. The fluorescence decay rates from the stopped flow measurements were compared with GLM using the Duncan’s Multiple Range Test.

## Acknowledgments

We thank Mikael Brosché for providing the seeds of the knockout mutants.

This work was funded by the Finnish Cultural Foundation.

